# Computational modeling of reinforcement learning and functional neuroimaging of probabilistic reversal dissociates compulsive behaviors in Gambling and Cocaine Use Disorders

**DOI:** 10.1101/2023.03.06.531272

**Authors:** Katharina Zühlsdorff, Juan Verdejo-Román, Luke Clark, Natalia Albein-Urios, Carles Soriano-Mas, Rudolf N. Cardinal, Trevor W. Robbins, Jeffrey W. Dalley, Antonio Verdejo-García, Jonathan W. Kanen

## Abstract

Cognitive flexibility refers to the ability to adjust to changes in the environment and is essential for adaptive behavior. It can be investigated using laboratory tests such as probabilistic reversal learning (PRL). In individuals with both Cocaine Use Disorder (CUD) and Gambling Disorder (GD), overall impairments in PRL flexibility are observed. However, it is poorly understood whether this impairment depends on the same brain mechanisms in cocaine and gambling addictions. Reinforcement learning (RL) is the process by which rewarding or punishing feedback from the environment is used to adjust behavior, to maximise reward and minimise punishment. Using RL models, a deeper mechanistic explanation of the latent processes underlying cognitive flexibility can be gained. Here, we report results from a re-analysis of PRL data from control participants (n=18) and individuals with either GD (n=18) or CUD (n=20) using a hierarchical Bayesian RL approach. We observed significantly reduced ‘stimulus stickiness’ (i.e., stimulus-bound perseveration) in GD, which may reflect increased exploratory behavior that is insensitive to outcomes. RL parameters were unaffected in CUD. We relate the behavioral findings to their underlying neural substrates through an analysis of task-based fMRI data. We report differences in tracking reward and punishment expected values (EV) in individuals with GD compared to controls, with greater activity during reward EV tracking in the cingulate gyrus and amygdala. In CUD, we observed reduced responses to positive punishment prediction errors (PPE) and increased activity following negative PPEs in the superior frontal gyrus compared to controls. Thus, an RL framework serves to differentiate behavior in a probabilistic learning paradigm in two compulsive disorders, GD and CUD.

## Introduction

The diagnostic criteria for both Substance Use Disorders (SUD) and Gambling Disorder (GD) in the Diagnostic and Statistical Manual of Mental Disorders (fifth edition) (DSM-5) include unsuccessful attempts to stop substance abuse or gambling, jeopardizing relationships and educational/career opportunities, and financial troubles arising as a consequence of the disorder (APA 2013). Compulsivity, a key feature of both GD and SUDs, is defined as persistent actions inappropriate to a given situation, which have no clear relationship to the overall goal and frequently result in undesirable consequences (Dalley et al. 2011). GD and SUDs are disorders of compulsivity and their behavioral phenotypes may thus overlap, but also diverge in certain aspects (Leeman and Potenza 2012; Robbins et al. 2012). Gaining a clearer definition of these phenotypes could inform the development of new treatments for disorders of compulsivity.

A further common feature of GD and SUD is behavioral inflexibility, defined as a deficit in adjusting behavior based on changes in environmental feedback (Jara-Rizzo et al. 2020; Smith et al. 2020; Perandrés-Gómez et al. 2021). Individuals with SUDs to a range of specific substances exhibit an increase in perseverative responding following a contingency change during probabilistic reversal learning (PRL), a paradigm used to investigate cognitive flexibility (Ersche et al. 2011). Increased perseveration during reversal is observed in individuals with Cocaine Use Disorder (CUD) (Ersche et al. 2008; Robinson et al. 2021). Indeed, reversal learning is impaired in rats and monkeys following prolonged exposure to cocaine (Jentsch et al. 2002; Schoenbaum et al. 2004).

Patients with GD, in comparison, show difficulties in learning novel stimulus-outcome associations following contingency changes during reversal learning (Jara-Rizzo et al. 2020). Following repeated negative feedback, patients with GD tend to stay rather than switch their response, or switch prematurely after little or no negative feedback during PRL (Perandrés-Gómez et al. 2021). Individuals with GD perform significantly worse than healthy controls (HCs) on the Intra-/Extra-Dimensional Set Shifting test (IED), which assays higher order cognitive flexibility, with impairments observed at the extra-dimensional shift stage (requiring the most flexibility) (Ornstein et al. 2000; Leppink et al. 2016). In a meta-analysis of nine studies that investigated performance of participants diagnosed with GD on the related Wisconsin Card Sorting Test (WCST), patients made more perseverative errors than healthy individuals (van Timmeren et al. 2018). Overall, it is evident that individuals with GD are impaired on cognitive flexibility tasks and have greater perseverative tendencies, similar to individuals with SUD.

CUD has been associated with fronto-striatal neuroadaptations that are linked to altered reward processing. For example, a study employing functional magnetic resonance imaging (fMRI) has found that individuals diagnosed with CUD exhibited lower blood-oxygen level dependent (BOLD) signals in the orbitofrontal cortex (OFC) than control participants following monetary gains on a forced-choice task containing three monetary value conditions (Goldstein et al. 2007). Neural activity is also known to be altered in patients with SUD during PRL, such as in the middle frontal gyrus (MFG) and caudate nucleus, areas known to contribute to performance on this task (Cools et al. 2002; Ersche et al. 2011). A meta-analysis of 52 studies reported that the OFC is hypoactive following detoxification in participants with CUD across different decision-making tasks (Dom et al. 2005). Thus, it is evident that activity of striatal and prefrontal cortical (PFC) regions is altered in CUD.

Functional MRI studies in individuals with GD have also found differential recruitment of PFC areas during reward-based tasks (Leeman and Potenza 2012). The ventromedial PFC (vmPFC), an area activated during monetary reward tasks in healthy individuals that is important for reward processing, shows decreased task-related activation in GD (Campbell-Meiklejohn et al. 2008; Habib and Dixon 2010; Li et al. 2010). On the Iowa Gambling Task, greater activity in individuals with GD during high-risk choices has been reported in the right caudate, OFC, vmPFC, superior frontal gyrus (SFG), amygdala, and hippocampus (Power et al. 2012). Furthermore, lower activity in the right ventrolateral PFC (vlPFC) has been linked to increased perseveration on a PRL task (De Ruiter et al. 2008). These findings point to altered reward processing in GD and suggest the involvement of cortical areas such as the vmPFC and OFC as well as subcortical structures; several areas overlap with those also affected in CUD.

Reinforcement learning (RL) is the process by which positive and negative feedback from the environment is used to adjust behavior to maximize rewards and minimize punishment (Sutton and Barto 1998). In recent years, RL models have increasingly been used to gain deeper insight into the latent mechanisms (represented by model parameters) underlying PRL. With RL models, for example, it is possible to interrogate how exploration versus exploitation (of learned values) contribute to choice behavior, or the degree of simple choice repetition unrelated to outcomes and value (stickiness). Reward and punishment learning rates can also be determined via RL models, which index the speed at which the expected value of a choice is updated after a better than or worse than expected outcome (reward or punishment prediction error). Indeed, RL impairments following drug use and withdrawal have been demonstrated in rodents and humans. In rats, increased exploitation and stickiness have been reported after cocaine self-administration, consistent with perseveration (Zhukovsky et al. 2019). Humans with SUD, meanwhile, have been found to be higher in stickiness and punishment learning rate, and have a lower reward learning rate (Kanen et al. 2019). RL modeling has also revealed that LSD, a non-selective serotonin 2A receptor agonist, increases reward and punishment learning rates and decreases stimulus stickiness in healthy volunteers (Kanen et al. 2022). Critically, the RL fingerprint during PRL in GD has not been elucidated. Furthermore, the neural substrates underlying these changes in RL parameters are not clearly defined. In rats, stickiness positively correlated with resting-state fMRI activity between the medial OFC (mOFC), PFC and subcortical structures (Zühlsdorff et al. 2023). In humans, the link between RL behavior and neural activity has not yet been established.

Here, we present a re-analysis of a previously published dataset (Verdejo-Garcia et al. 2015) using novel computational methods. Individuals with CUD, GD and controls completed a PRL task in an fMRI scanner. In the previous publication arising from this dataset, conventional PRL measures were calculated and compared between the groups. There, it was reported that a behavioral variable reflecting the perseveration error rate was increased in GD, with no changes observed in the CUD group. Additionally, both patient groups had lower vlPFC activation when shifting responding following a reversal. When perseverating, participants with CUD had greater activity in the dorsomedial PFC (dmPFC) than the GD group. In the new analysis presented here, RL models are employed to reveal latent processes underlying behavior on the PRL task, via a potentially more sensitive trial-by-trial approach. Through the fMRI data, the RL parameters can be linked to their associated neural substrates. To our knowledge, no previous studies have investigated PRL data from GD patients using RL models. Based on our recent work that showed the concept of stickiness (the tendency to repeat actions regardless of value) was critical for dissociating other disorders of compulsivity (Kanen et al. 2019), we hypothesized that stickiness would be central to the RL modeling results here and would be increased in GD and CUD. Neurally, we predicted that activity in the amygdala and OFC would be linked to the reward learning rate, and that medial PFC and dorsal striatal activity would reflect the stickiness parameter.

## Methods

### Participants

Fifty-six participants took part in this study. These comprised: 18 healthy control subjects who did not meet any of the criteria for an Axis I or II disorder; 18 subjects who met the DSM-IV-TR criteria for Gambling Disorder and 20 individuals that met the criteria for Cocaine Use Disorder. Here, we use the term Substance Use Disorder as used in the DSM-V, rather than stimulant dependence, which is used in the DSM-IV-TR (APA 2013). Basic behavioral data in association with fMRI findings from this study have previously been published (Verdejo-Garcia et al. 2015).

Individuals with CUD were recruited in the outpatient clinic Centro Provincial de Drogodependencias, Granada, Spain, and participants diagnosed with GD were recruited from Asociación Granadina de Jugadores en Rehabilitación, Granada, Spain. Participants from these two groups had met the following inclusion criteria: 1) between 18-45 years old; 2) estimated IQ level above 80; 3) meeting the DSM-IV-TR criteria for cocaine dependence or pathological gambling; 4) having commenced psychological treatment; 5) having been abstinent for more than 15 days. It was confirmed that individuals with CUD were abstinent using two urine tests per week and an additional test on the imaging day. Gambling abstinence was confirmed by relatives and checked through self-assessment. The following exclusion criteria were applied: 1) diagnosis of another Axis I or II disorder, except alcohol or nicotine addiction; 2) history of head injury, neurological disease or any other diseases affecting the central nervous system; 3) having undertaken other treatments in the two years prior to the study; 4) court-mandated treatment. The Structured Clinical Interview for DSM-IV Axis I Disorders (SCID-I-CV) was used to assess Axis I disorders, whereas Axis II disorders were assessed through the International Personality Disorders Examination (IPDE) (Loranger 1994; First 1997). Diagnoses were made through registered clinical psychologists. Control participants were recruited from local agencies. The study was approved by the Ethics Committee for Research in Humans, University of Granada, Spain. Participants signed an informed consent form to confirm their voluntary participation and were all equally reimbursed for their participation.

### Probabilistic Reversal Learning Task

This task was similar to the PRL task used by (Cools et al. 2002). Two abstract, colored stimuli were presented on the right and left side of the visual display. Stimulus location was randomized. At the beginning of the tasks, everyone was informed that one stimulus was the ‘correct’ stimulus (CS+), and the other stimulus was the ‘incorrect’ stimulus (CS−). Subjects had to learn the correct and incorrect stimulus through a trial-and-error approach. The CS+ resulted in a reward on only 85% of the trials, whereas the CS− was rewarded 15% of the time. Following 10 to 15 correct trials, the contingencies were reversed. All participants were trained on the PRL task outside the scanner before the initial scan, for which different stimuli were used. During scanning, there were three consecutive blocks that consisted of 10 discriminations (9 reversals), with a duration of 11 min per block.

Magnetic-resonance-compatible liquid-crystal display goggles were used to present the stimuli (Resonance Technology Inc., Northridge, CA, USA). All responses were recorded using the Evoke Response Pad System (Resonance Technology Inc.). This button box was located on the subject’s chest. The duration of stimulus presentation was 2000 ms. If participants failed to respond during this time, a ‘too late’ message was presented. Following a ‘correct’ response, a green smiley face was presented, and following an ‘incorrect’ response, a red sad face was shown. Feedback was presented for 500 ms, during which time the stimulus remained on the screen. Following feedback presentation, there was a variable inter-trial interval, which was adjusted by the program, for a final interstimulus interval duration between stimuli of 3253 ms. This interstimulus interval duration was selected to enable a precise desynchronization from the repetition time (2000 ms).

### Reinforcement learning modeling

The PRL data was modelled with RL models using a hierarchical Bayesian approach. Seven different models containing different combinations of RL parameters (described in more detail below) were tested, implemented through Stan (Stan Development Team 2020).

The highest hierarchical level contained a group-specific mean and a common standard deviation for every RL parameter of the respective model. Priors for these values are shown in **Table 1**. The RL parameters were drawn for each subject from a normal distribution having the relevant mean/standard deviation. Predicted choices were fit to behavior according to an RL algorithm (described below), and the highest posterior density interval (HDI) was calculated for group mean differences of interest (Kruschke 2014).

**Table 1.** Priors for model parameters.

| Parameter | Prior | Reference |
| --- | --- | --- |
| Reward learning rate, $\alpha_{rew}$ | Beta(1.2, 1.2) | (Den Ouden et al. 2013) |
| Punishment learning rate, $\alpha_{pun}$ | Beta(1.2, 1.2) | (Den Ouden et al. 2013) |
| Combined learning rate, $\alpha$ | Beta(1.2, 1.2) | (Den Ouden et al. 2013) |
| Reinforcement sensitivity, $\beta$ | Gamma($\alpha=4.82$, $\beta=0.88$) | (Gershman 2016) |
| Side stickiness, $\kappa_{side}$ | Normal(0,1) | (Christakou et al. 2013) |
| Stimulus stickiness, $\kappa_{stim}$ | Normal(0,1) | (Christakou et al. 2013) |
| Experience decay factor, $\rho$ | Beta(1.2, 1.2) | (Den Ouden et al. 2013) |
| Decay factor for previous payoffs, $\phi$ | Beta(1.2, 1.2) | (Den Ouden et al. 2013) |
| Softmax inverse temperature, $\beta$ | Gamma($\alpha=4.82$, $\beta=0.88$) | (Gershman 2016) |
| **Intersubject variability in parameters** |  |  |
| $\alpha_{rew}$, $\alpha_{pun}$, $\alpha$, $\kappa_{side}$, $\kappa_{stim}$, $\rho$, $\phi$ intersubject standard deviations | Normal(0,0.05) constrained to $\geq 0$ | (Christakou et al. 2013; Den Ouden et al. 2013) |
| $\beta$ intersubject standard deviations | Normal(0,1) constrained to $\geq 0$ | (Gershman 2016) |

Q values were updated on a trial-by-trial basis according to the following equation:

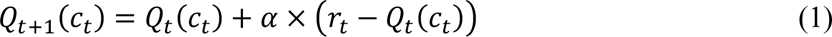

*Q_t_*_+1_(*ct*) is the expected value for the next trial based on the stimulus that is chosen on the current trial, *Q_t_*(*c_t_*) is the expected value of the choice taken on the current trial, *α* is the learning rate and *r_t_* is the reinforcement on trial *t* (1 for reward and 0 for punishment). The learning rate influences how much the subject updates the Q value based on the prediction error *r_t_* − *Qt*(*ct*), with higher α driving faster learning.

The probability of making one of two choices given the Q values for each was calculated using the softmax decision rule:

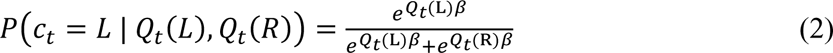

Q_t_(L) and Q_t_(R) are the Q values of the left and right stimuli, and β is the reinforcement sensitivity parameter, which determines to what extent the subject is driven by its reinforcement history (versus random choice).

The seven models tested were as follows:

1. **Two parameters: α and β**, the learning rate and reinforcement sensitivity parameter.
2. **Three parameters: α, β,** stimulus stickiness parameter **κ_stim_**. κ_stim_ is the tendency to respond to the same stimulus as on the previous trial, irrespective of its location and outcome (i.e., whether it was rewarded or not), and was used to update the Q value as follows: *Q*^*stim*^ *_s,t_*_+1_= κ_*stim*_*S*_*s*,*t*_. *S*_*s*,*t*_ represents the stimulus chosen by the subject on the last trial. This value is 1 if the same stimulus was chosen, and 0 if another stimulus was chosen. The final Q value is the sum of *Q*^*stim*^_*s,t*+1_ and the Q value as calculated in equation 1.
3. **Three parameters: α_rew_, α_non-rew_, β.** Similar to model 1, but containing two separate learning rates for rewarded and non-rewarded trials, respectively.
4. **Four parameters: α_rew_, α_non-rew_, β and κ_stim_.**
5. **Four parameters: α_rew_, α_non-rew_, β and κ_side_.** The stimulus stickiness parameter was replaced with the side stickiness parameter, representing the tendency to choose the same side as on the previous trial, irrespective of the outcome produced, and was used to update the Q value as follows: *Q*^*loc*^_*s*,*t*+1_ = κ_*side*_ *L*_*s*,*t*_. *L*_*s*,*t*_ represents the side chosen by the subject on the last trial. This value is 1 if the same side was chosen, and 0 if the other side was chosen. The final Q value is the sum of *Q*^*loc*^_*s*,*t*+1_ and the Q value as calculated in equation 1.
6. **Five parameters: α_rew_, α_non-rew_, β, κ_stim_ and κ_side_.**
7. **The Experience-Weighted Attractor model (EWA) (Camerer and Ho 1999).**

In this model, the value of incoming information is compared against the individual’s beliefs. The parameters included are experience weight φ for each stimulus, which modulates learning from reinforcement. This value changes over time according to the decay factor ρ. This model also includes the parameter **β**.

The models were fitted through Hamiltonian Markov Chain Monte Carlo sampling via Stan 2.17.2 (Carpenter et al. 2017). Convergence was ensured using the potential scale reduction factor (Brooks and Gelman 1998; Gelman 2013). A potential scale reduction factor value close to 1 indicated perfect convergence. A cut-off of 1.1 was selected as a stringent criterion for convergence. Models were compared using a bridge sampling estimate of the marginal likelihood using the “bridgesampling” R package (Gronau et al. 2017, 2020).

Between-group differences were sampled to give a posterior probability distribution for each quantity of interest. These posterior distributions were interpreted using the 95% and 75% HDI, which are ‘credible intervals’ in Bayesian statistics. At 95% HDI, more evidence is provided for there being group differences than at 75% HDI. However, findings at 75% HDI are also considered to provide sufficient evidence for there being group differences.

### Data simulation

Data were simulated using the posterior group mean parameters from the winning model, with the aim of determining whether the winning model could reproduce the behavioral observations. The simulated data were then analyzed using a conventional PRL analysis as described in (Verdejo-Garcia et al. 2015). One hundred virtual “subjects” were simulated for each group, with each “subject” performing the PRL task in silico.

### Imaging acquisition

Subjects were scanned in a 3T MRI scanner with an eight-channel phased-array head coil (Intera Achieva, Philips Medical Systems, Eindhoven, The Netherlands). First, three T2*-weighted scans using an echo planar imaging (EPI) sequence were taken (repetition time (TR)=2000 ms, time to echo (TE)=35 ms, field of view (FOV)=230×230 mm, 96×96 matrix, flip angle=90°, 21 4-mm axial slices, 1-mm gap, 330 scans each). Subsequently, a sagittal three-dimensional T1-weighted turbo-gradient-echo sequence was used (150 slices, TR=8.3 ms, TE=3.8 ms, flip angle=8°, FOV=240×240, 1 mm^3^ voxels). More details can be found in (Verdejo-Garcia et al. 2015).

### Image pre-processing

The FMRIB Software Library (FSL) and FMRIPREP were used to pre-process the data (Smith et al. 2004; Esteban et al. 2018). FMRIPREP implements multiple software, including FSL and the Advanced Normalisation Tools (ANTs) (Tustison et al. 2010). Each T1-weighted image was bias-field corrected using *N4BiasFieldCorrection* and skull-stripped using *antsBrainExtraction* with the OASIS template from the ANTs software. Functional MRI scans were spatially normalized to the ICBM 152 Nonlinear Asymmetrical template version 2009c through non-linear registration with the *antsRegistration* tool using brain-extracted versions of both the T1-weighted (T1w) volume and template (Avants et al. 2008).

Subsequently, brain extracted T1w images were segmented into cerebrospinal fluid, white matter and grey matter using *fast* (FSL) (Zhang et al. 2001). Functional MRI scans were slice-timing-corrected using *slicetimer* (FSL) and then motion-corrected with *mcflirt* (FSL) (Jenkinson et al. 2002). For scans with associated field maps, distortion correction was performed using *fugue* (FSL) (Jenkinson 2003). Next, the fMRI images were co-registered to their corresponding T1w scan using boundary-based registrations with six degrees of freedom with *flirt* (FSL) (Greve and Fischl 2009). The field distortion correcting warp, BOLD-to-T1w transformation and T1w-to-template (MNI) warp were concatenated and applied in a single step using *antsApplyTransforms* using Lanczos interpolation. Nipype was used to calculate the frame-wise displacement (Power et al. 2014). The first five volumes were discarded to avoid T1 saturation effects. fMRI images were high-pass filtered (128 s) and spatially smoothed with a 6 mm full-width, half-maximum 3D Gaussian kernel. A canonical hemodynamic response function was modelled to the onsets of the explanatory event types. Multiple criteria were used to ensure successful registration, including checking successful registration, ensuring that none of the participants showed excessive motion using DVARS (root mean square of the temporal change of the voxel-wise signal at each time point (Yang et al. 2019)) and framewise-displacement measures (excessive motion threshold being 10% of the total number of volumes) and by inspecting their respective carpet plots.

### First-level models

First-level linear models were fit through FEAT (FSL) (Woolrich et al. 2001). A first-level model was fit for each run and included the following event types: (1) reward Expected Value (EV), (2) positive Reward Prediction Error (RPE), (3) negative RPE, (3) punishment EV, (5) positive Punishment Prediction Error (PPE), (6) negative PPE and (7) response/feedback presentation. The RPE is representative of a predicted reward and is positive when there is an unexpected or better than expected reward, and negative if an expected reward is omitted or the outcome is worse than expected. The PPE is when a punishment is expected. Similarly to the RPE, it is positive if a reward is received, and negative if there is a punishment or no reward. EV and prediction errors (PEs) were extracted for each trial from the winning Q-learning model. Explanatory variables 1–6 were based on the extracted values of prediction error and reward or punishment cue values. Positive PEs took values between 0 and 1, whereas negative PEs were between 0 and −1. The model was based on an analysis presented previously (Murray et al. 2019). Six movement parameters (x, y, z, pitch, roll, yaw) were incorporated into the model, which resulted from the image realignment to control for movement artefacts.

### Higher-level models

The first-level models were averaged across the three runs for each subject, resulting in the second-level models. Third-level mixed-effects whole-brain analyses involving one-sample t-tests with cluster thresholding with a Z threshold of ±3.1 and p<0.05 were used to investigate the contrasts for each event type (Woolrich et al. 2004). The contrasts included control vs GD, control vs CUD and GD vs CUD. Subsequently, an analysis of covariance (ANCOVA) was run as an additional exploratory analysis. In the ANCOVA, model parameters from the best-fitting RL model were extracted for each subject and included as predictors. The aim of this analysis was to investigate group differences in the correlation between activity in a given region and a RL parameter (i.e., a group×RL parameter interaction). RL parameters were also correlated with BOLD signal from all participants, regardless of group. FSLeyes was used to generate figures (Smith et al. 2004). In all figures, the right and left sides are inverted from the observer’s perspective (according to standard radiological convention).

## Results

### Demographic information

There were no significant differences in age, gender, IQ, handedness, or years of education between the groups (**Table 2**) (Verdejo-Garcia et al. 2015).

**Table 2.** Demographic information.

|  | **Healthy Controls (n=18)** | **Gambling Disorder (n=18)** | **Cocaine Use Disorder (n=20)** | **Group Comparisons** |
| --- | --- | --- | --- | --- |
| Mean age (SD) | 31.2 (4.7) | 33.6 (8.0) | 34.3 (6.9) | F(2,54)=1.43, p=0.35 |
| Gender (Female) | 1 | 2 | 1 | $X^2(2,56)=0.59$, p=0.75 |
| Verbal IQ (SD) | 106.9 (9.0) | 102.7 (7.4) | 100.9 (7.6) | F(2,54)=2.31, p=0.082 |
| Years of Education (SD) | 10.6 (1.9) | 10.3 (2.1) | 9.8 (1.7) | F(2,54)=1.37, p=0.47 |
| Handedness (L) | 1 | 1 | 4 | $X^2(2,56)=2.80$, p=0.25 |

L, left; SD, standard deviation.

### Selecting the winning model

**Table 3** reports the results from the seven RL models tested and model comparison measures. Satisfactory model convergence was confirmed, as all parameters and contrasts had a potential scale reduction factor of less than 1.1, with the maximum value being 1.006.

**Table 3.** Model comparison summary. Models were assumed to be equiprobable *a priori*.

| **Model** | **Rank** | **Parameters** | **Log marginal likelihood** | **Log posterior P** |
| --- | --- | --- | --- | --- |
| 5 | 7 | $\alpha_{rew}$, $\alpha_{non-rew}$, $\beta$, $\kappa_{side}$ | -13168.48 | -582.60 |
| 6 | 1 | $\alpha_{rew}$, $\alpha_{non-rew}$, $\beta$, $\kappa_{side}$, $\kappa_{stim}$ | -11139.35 | 0.000 |
| 4 | 2 | $\alpha_{\text{rew}}, \alpha_{\text{non-rew}}, \beta, \kappa_{\text{stim}}$ | -13003.72 | -417.85 |
| 3 | 4 | $\alpha_{\text{rew}}, \alpha_{\text{non-rew}}, \beta$ | -13130.08 | -544.21 |
| 2 | 6 | $\alpha, \beta, \kappa_{\text{stim}}$ | -13018.42 | -432.54 |
| 7 | 3 | $\phi, \rho, \beta$ | -13161.24 | -575.36 |
| 1 | 5 | $\alpha, \beta$ | -13159.06 | -573.18 |

The winning model (model 2) contained five parameters: the reward learning rate *α*_*rew*_, representative of how quickly an individual updates (increases) Q values in response to positive feedback; the punishment learning rate *α*_*pun*_, reflecting how quickly an individual updates (decreases) the Q-value following punishment; reinforcement sensitivity *β*, also known as the exploitation vs exploration or inverse temperature parameter; stimulus stickiness ^κ^*stim*, which is the tendency to select the same stimulus regardless of outcome, and side stickiness ^κ^*side*, the tendency to select the same side regardless of outcome.

### Reinforcement learning results

**Figure 1** shows results of the hierarchical Bayesian RL analysis. Neither the reward learning rate nor the punishment learning rate were affected in GD or CUD when compared with healthy controls. However, there was evidence that the reward learning rate *α*_*rew*_ was lower in the CUD group than the GD group (difference in parameter per-group mean, posterior 75% HDI excluding zero). Reinforcement sensitivity was lower in the CUD group compared to the GD group, reflecting more exploratory behavior in CUD (group difference, 0 ∉ 75% HDI). Side stickiness, meanwhile, was not different in either patient group compared to the control group (no group differences, 0 ∈ 75% HDI). There was evidence for a decrease in stimulus stickiness at 75% HDI in the GD group compared to HCs (group difference, 0 ∉ 75% HDI). There were no changes in the CUD group when compared to the control group (no group differences, 0 ∈ 75% HDI). To summarize, we found evidence for the stimulus stickiness parameter ^κ^*stim* being decreased in the GD group. Moreover, when comparing the GD group to the CUD group, there was support for the reward learning rate *α*_*rew*_ and reinforcement sensitivity parameter *β* being greater in the GD group compared to the CUD group. No differences at 95% HDI were observed.

**Figure 1.**
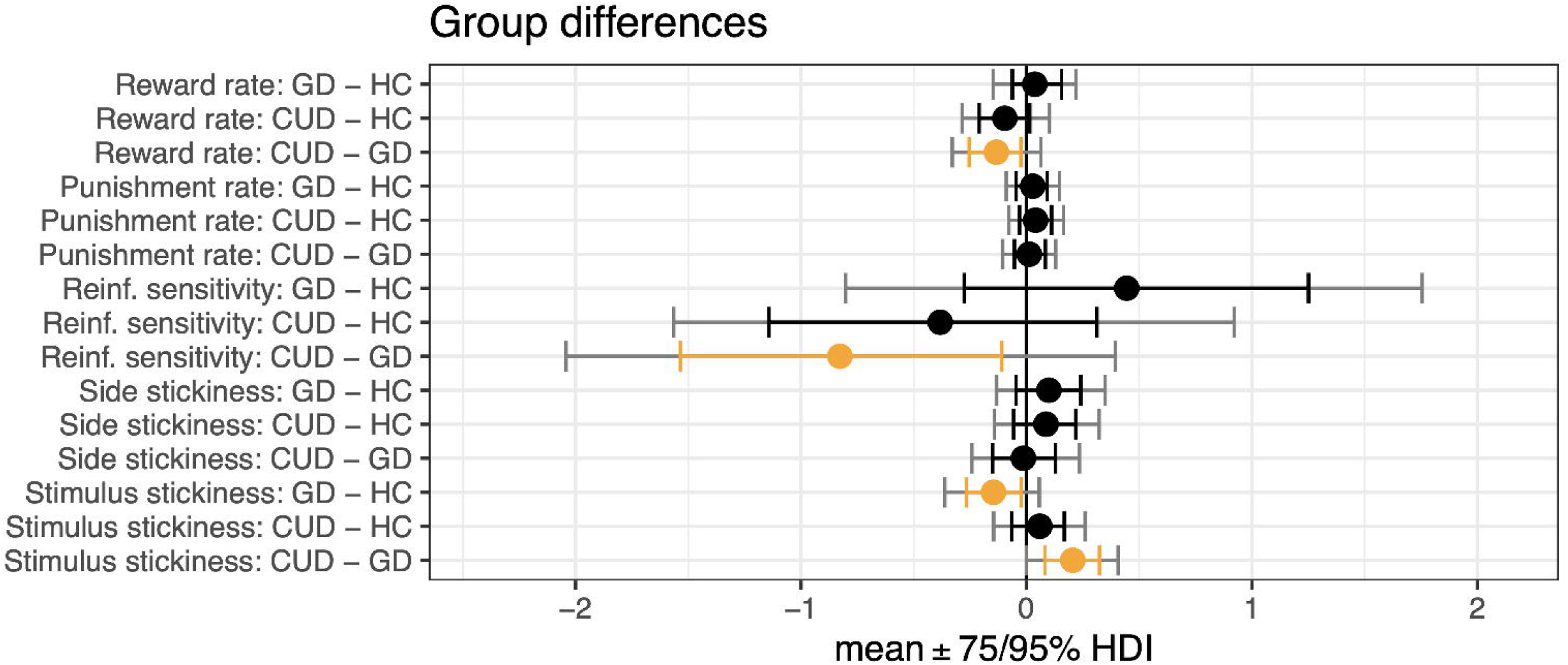
Results from the hierarchical Bayesian winning RL model, showing differences in group mean parameters. GD, Gambling Disorder; CUD, Cocaine Use Disorder; HC, healthy controls; Reinf. sens, reinforcement sensitivity; Stim, stimulus; HDI, highest posterior density interval. Orange indicates 75% HDI.

### Simulations

The parameters from the winning RL model were used to simulate the behavioral data and determine whether this model could replicate the behavior observed initially via raw data measures. When these data were analyzed using a conventional approach to extract raw data measures such as win-stay and lose-shift, no statistically significant differences between the groups were found. These findings thus align with the results for the conventional behavioral measures presented in (Verdejo-Garcia et al. 2015), suggesting that the model was able to reproduce the behavioral dynamics on this task.

### Brain activity during reward and punishment expected value tracking in Gambling Disorder

The model fitted to the task-based fMRI data included seven explanatory variables, as above: (1) reward EV; (2) positive RPE; (3) negative RPE; (4) punishment EV; (5) positive PPE; (6) negative PPE and (7) response/feedback presentation. We found differences in the neural responses to reward and punishment expected value in the GD group compared to controls. Specifically, we observed that when tracking reward EV, that individuals with GD had greater activations in the amygdala, hippocampus, parahippocampal gyrus, lateral occipital cortex, superior, inferior, and middle temporal gyri, as well as the precuneus than HCs (**Figure 2**, **Table 4**). These effects were only observed in the left hemisphere.

**Figure 2.**
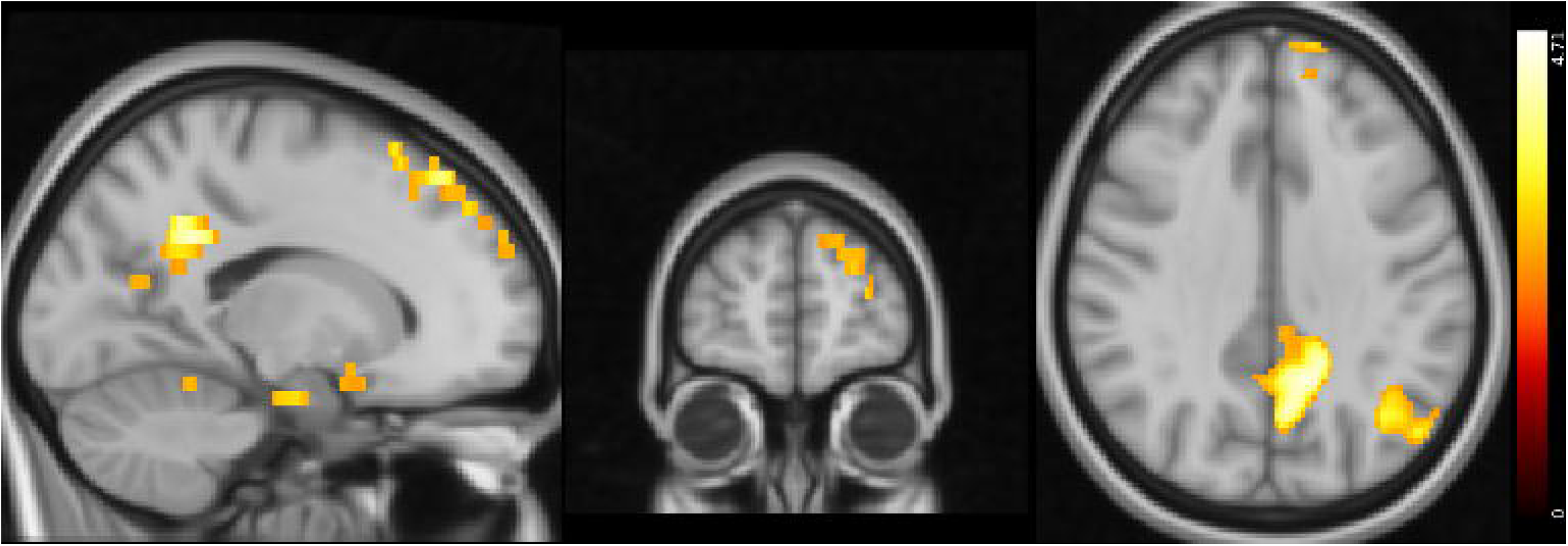
Reward EV tracking: differences between healthy controls and participants with GD (MNI coordinates: X=-16, Y=58, Z=34). Activity was higher in the GD group in the areas indicated. Color bar on the right-hand side represents the *t* statistic.

**Table 4.**
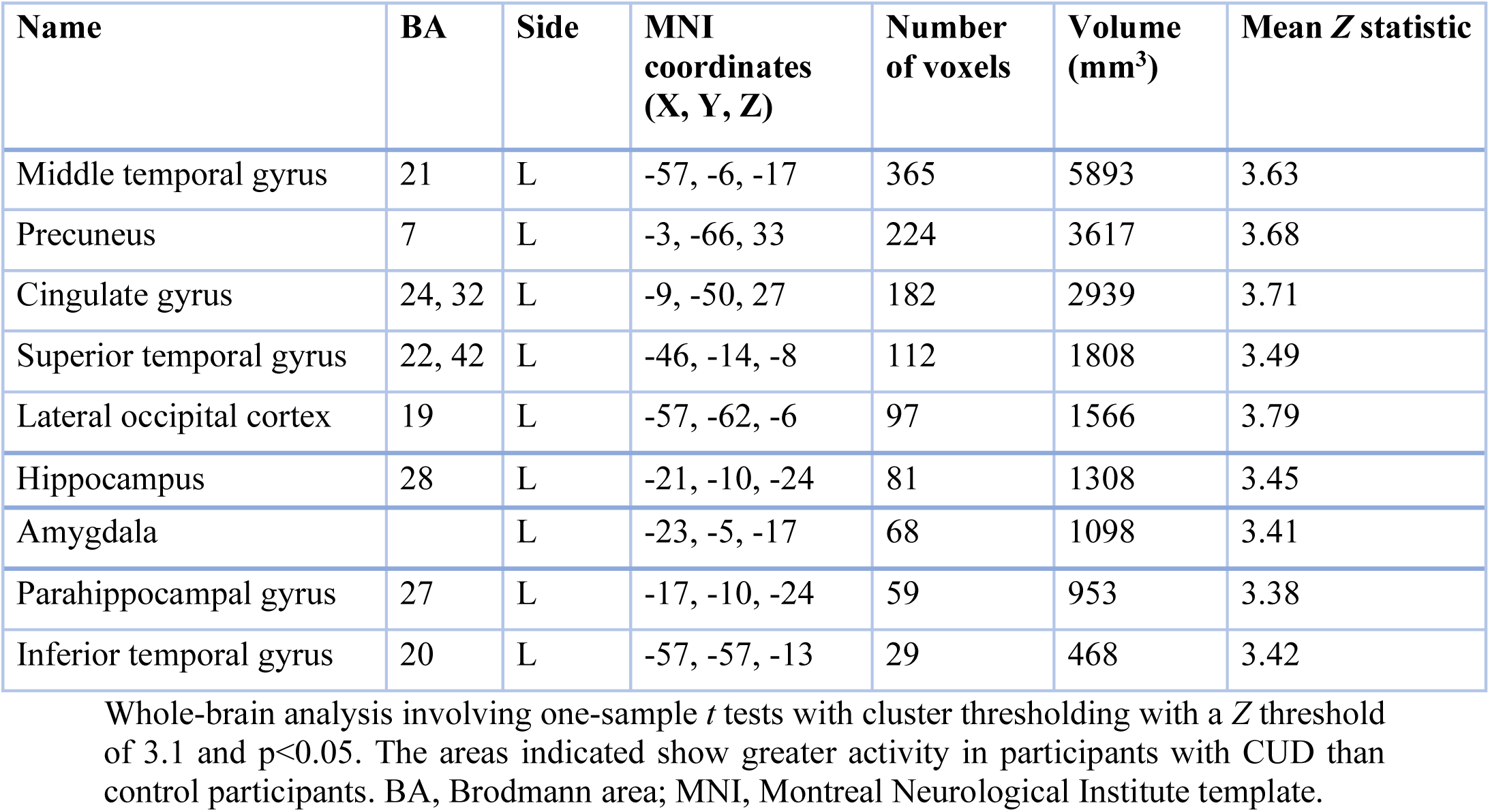
Summary of peak fMRI activity for the reward EV controls-vs-GD contrast.

For punishment EV, we observed the opposite trend: individuals with GD showed lower activity in the superior parietal lobule, pre- and postcentral gyri, precuneus, parietal operculum, supramarginal gyrus and angular gyrus compared to control subjects (**Figure 3**, **Table 5**). Activations were seen in both hemispheres but were more pronounced in the right hemisphere.

**Figure 3.**
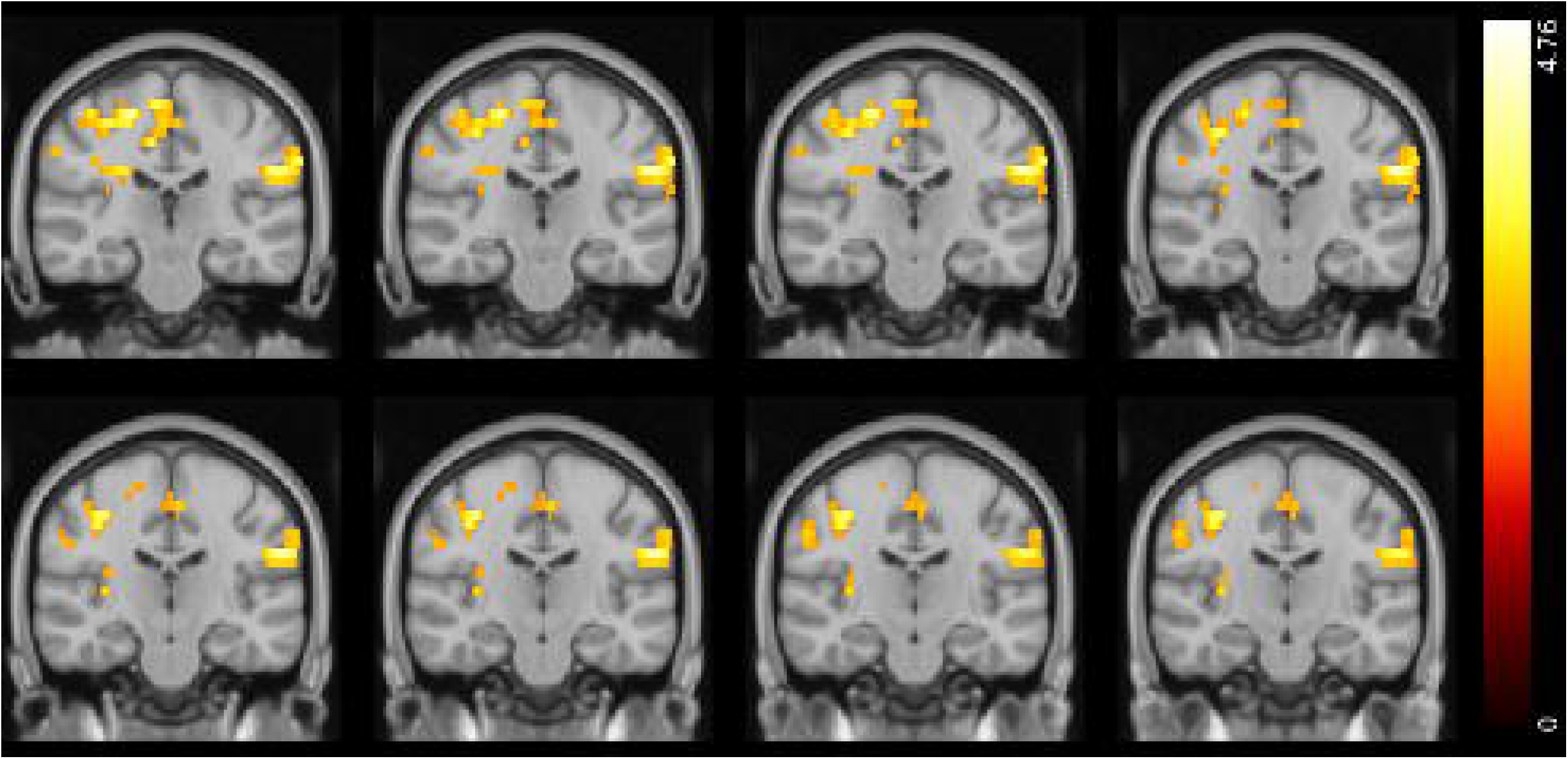
Punishment EV tracking: differences between healthy controls and participants with GD (MNI coordinates: Y=-24 to −17). Activity was lower in the GD group in the areas indicated. Color bar on the right-hand side represents *t*.

**Table 5.** Summary of peak fMRI activity for the punishment EV controls-vs-GD contrast.

| Name | BA | Side | MNI coordinates (X, Y, Z) | Number of voxels | Volume (mm<sup>3</sup>) | Mean z-statistic |
| --- | --- | --- | --- | --- | --- | --- |
| Postcentral gyrus | 1, 2, 3 | R | 56, -14, -33 | 1228 | 19827 | 2.99 |
| Postcentral gyrus | 1, 2, 3 | L | -62, -21, -33 | 408 | 6588 | 3.02 |
| Precentral gyrus | 4 | R | 43, -14, 45 | 865 | 13966 | 2.95 |
| Precuneus | 7 | R | 10, -51, 56 | 555 | 8961 | 2.90 |
| Precuneus | 7 | L | -6, -48, 56 | 403 | 6507 | 2.90 |
| Superior parietal lobule | 7 | R | 28, -44, 59 | 524 | 8461 | 3.04 |
| Supramarginal gyrus | 40 | R | 56, -18, 31 | 254 | 4101 | 2.88 |
| Supramarginal gyrus | 40 | L | -63, -26, 31 | 258 | 4166 | 3.02 |
| Lateral occipital cortex | 19 | R | 17, -79, 41 | 242 | 3907 | 2.88 |
| Lateral occipital cortex | 19 | L | -14, -83, 41 | 181 | 2922 | 2.75 |
| Parietal operculum cortex | 40, 43 | R | 1.5, -33, 21 | 106 | 1711 | 2.83 |
| Parietal operculum cortex | 40, 43 | L | -51, -33, 21 | 191 | 3084 | 2.98 |

Whole-brain analysis involving one-sample $t$ tests with cluster thresholding with a $Z$ threshold of 3.1 and $p < 0.05$ . The areas indicated show lower activity in participants with CUD than control participants. BA, Brodmann area; MNI, Montreal Neurological Institute template.

### Neural signal to positive and negative punishment prediction errors is altered in Cocaine Use Disorder

We observed aberrant neural responses in CUD as well, specifically in response to positive and negative PPEs. Compared to control participants, individuals with CUD exhibited lower activity in the paracingulate gyrus and left SFG in response to positive PPEs. Conversely, individuals with CUD showed greater activity in the left SFG and MFG in response to negative PPEs (**Figures 4, 5**; **Tables 6, 7**, respectively).

**Figure 4.**
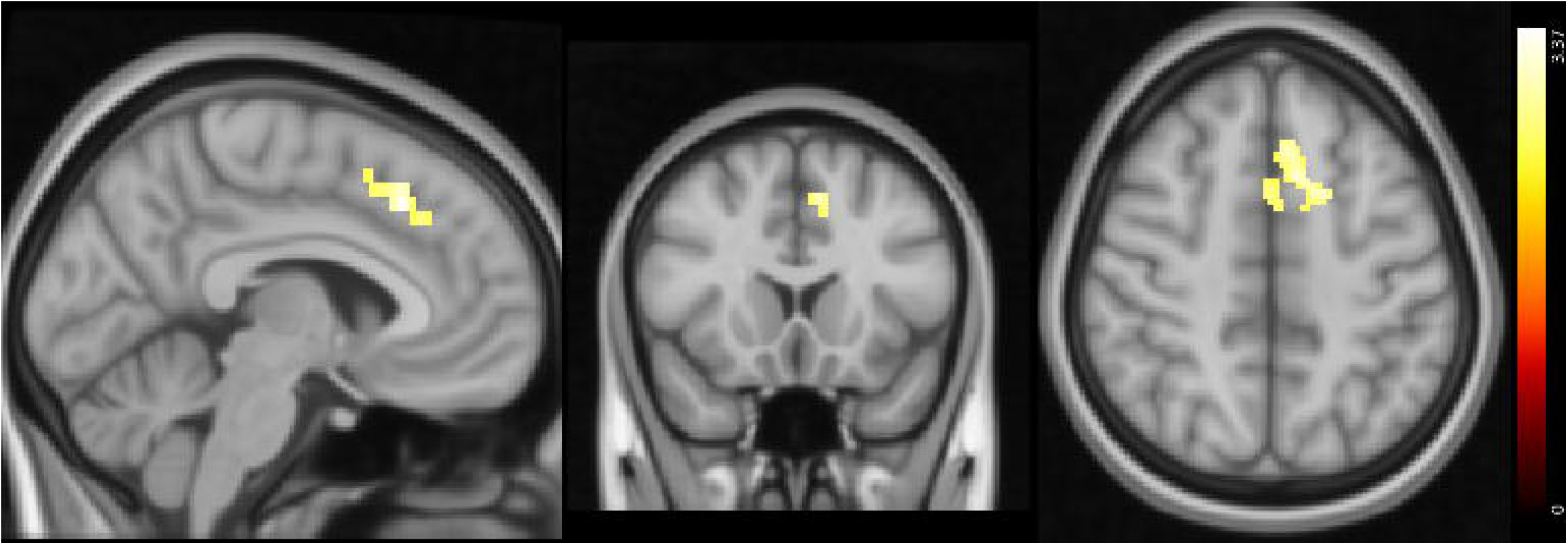
Response to positive PPE: differences between healthy controls and participants with CUD (MNI coordinates: X=-5, Y=17, Z=48). Activity was lower in the CUD group in the areas indicated. Color bar on the right-hand side represents *t*.

**Figure 5.**
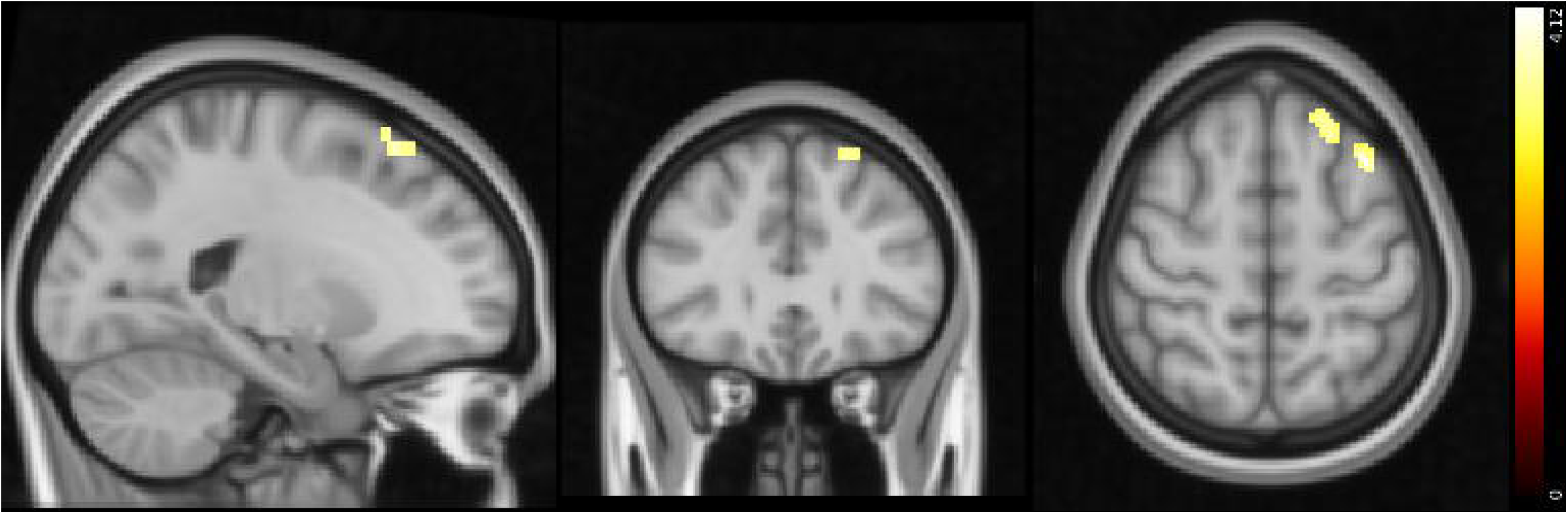
Response to positive PPE: differences between healthy controls and participants with CUD (MNI coordinates: X=-31, Y=30, Z=56). Activity was higher in the CUD group in the areas indicated. Color bar on the right-hand side represents *t*.

**Table 6.** Summary of peak fMRI activity for the positive PPE controls-vs-CUD contrast.

| Name | BA | Side | MNI coordinates (X, Y, Z) | Number of voxels | Volume (mm<sup>3</sup>) | Mean z-statistic |
| --- | --- | --- | --- | --- | --- | --- |
| Superior frontal gyrus | 8, 9 | L | -10, 13, 53 | 149 | 2406 | 2.81 |
| Paracingulate gyrus | 32 | L | -8, 20, 43 | 132 | 2131 | 2.80 |
| Paracingulate gyrus | 32 | R | 4, 11, 47 | 36 | 581 | 2.71 |

Whole-brain analysis involving one-sample $t$ tests with cluster thresholding with a $Z$ threshold of 3.1 and $p < 0.05$ . The areas indicated show lower activity in participants with CUD than control participants. BA, Brodmann area; MNI, Montreal Neurological Institute template.

**Table 7.** Summary of peak fMRI activity for the punishment PPE controls-vs-CUD contrast.

| Name | BA | Side | MNI coordinates (X, Y, Z) | Number of voxels | Volume (mm<sup>3</sup>) | Mean z-statistic |
| --- | --- | --- | --- | --- | --- | --- |
| Superior frontal gyrus | 8, 9 | L | -57, -6, -17 | 71 | 1146 | 3.41 |
| Middle frontal gyrus | 8, 9 | L | -3, -66, 33 | 70 | 1130 | 3.41 |

Whole-brain analysis involving one-sample $t$ tests with cluster thresholding with a $Z$ threshold of 3.1 and $p < 0.05$ . The areas indicated show greater activity in participants with CUD than control participants. BA, Brodmann area; MNI, Montreal Neurological Institute template.

### Neural responses to feedback presentation

During feedback presentation, the GD group overall showed increased activity (versus controls) in the lateral occipital cortex, cingulate gyrus, parahippocampal gyrus, precuneus, middle temporal gyrus and supramarginal gyrus (supplementary materials, **Figure S1**). There were also significantly greater activations during simultaneous cue and feedback presentation in the CUD group (versus controls), which were instead in the frontal pole, SFG, inferior frontal gyrus (IFG), precentral gyrus, superior parietal lobule, supramarginal gyrus, precuneus, angular gyrus, and lateral occipital cortex (**Figure S2**). Moreover, we observed differences between the CUD and GD groups: individuals with CUD had greater activity than those with GD in the insular cortex, IFG, and frontal operculum.

No significant differences were found in response to positive and negative RPEs. Thus, there appear to be widespread differences in both CUD and GD groups when the feedback was presented. However, this response was altered in different areas of the brain in the two disorder groups.

### Whole-brain correlation analyses

The five parameters from the winning RL model were used in a whole-brain correlation analysis to identify whether they correlated with the BOLD signal during each event type in any of the brain regions. This was done to identify the brain regions underlying RL parameters. The first analysis related the parameters to activity from all subjects.

This analysis highlighted that the *α*_*rew*_ parameter correlated negatively with activity in the cingulate and paracingulate gyri, IFG, middle and superior temporal gyri, insular cortex, and mOFC during reward EV tracking as well as responses to positive PPEs. This parameter also correlated negatively with activity in the putamen, mOFC, and insula during positive RPEs (see **Supplementary Materials**).

Next, an ANCOVA was run to compare connectivity patterns among the different groups. In the GD group, *α*_*rew*_ correlated more strongly with activity in the SFG, MFG, postcentral gyrus during reward EV tracking compared to the other two groups (**Figure S3**). In the CUD group, the correlation between *α*_*rew*_ and activity during the positive PPE was greater in the frontal pole, SFG, cingulate and paracingulate gyri compared to the HC and GD groups (**Figure S4**).

In both patient groups, stimulus stickiness (^κ^*stim*) had a stronger positive correlation with activity in the right MFG and IFG during cue/feedback presentation compared to control participants, suggesting that there are increased activations in these areas in patients when repeating a response regardless of previous outcomes (**Figure 6**). No other correlations with RL parameters were found.

**Figure 6.**
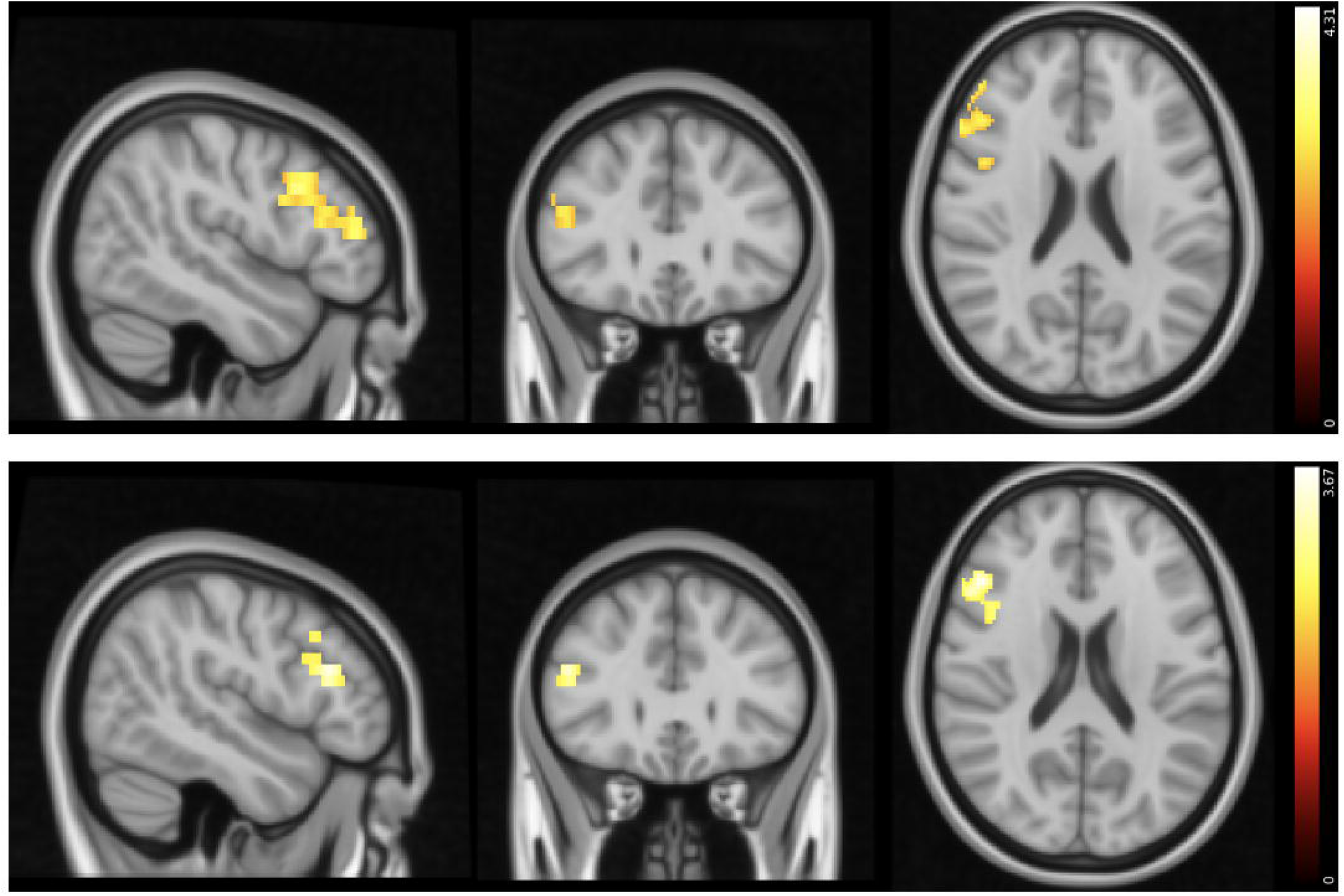
Top: Areas that have a stronger positive correlation with κ_stim_ in the GD group than in healthy controls (MNI coordinates: X=48, Y=29, Z=22). Bottom: Areas that have a stronger positive correlation with κ_stim_ in the CUD group than in healthy controls (MNI coordinates: X=48, Y=29, Z=20). Color bar on the right-hand side represents *t*.

## Discussion

In this study, we examined RL processes during a classic test of behavioral flexibility (PRL) in individuals with GD and CUD. Our computational modeling approach enabled the assessment of how both value-based (learning rates, reinforcement sensitivity) and value-free (stimulus and side stickiness) contributed to choice behavior. The key behavioral result was that individuals with GD showed reduced choice repetition (stimulus stickiness), irrespective of the feedback received, suggestive of a maladaptive exploratory pattern. Reduced stimulus stickiness in GD contrasts with our recent observation of abnormally increased choice repetition in SUD, regardless of reinforcement (Kanen et al. 2019). Stimulus stickiness (a form of choice repetition) may therefore present a novel way of dissociating compulsive disorders, in this case GD and CUD. However, we note that group differences were only observed at 75% HDI, but not at 95%.

We provide a novel and unexpected insight into how RL parameters are affected in GD – that stimulus stickiness was reduced in this group. A similar reduction in stimulus stickiness has also been observed in another compulsive disorder, OCD (Kanen et al. 2019). However, in GD, the reduction in stimulus stickiness was accompanied by a slight increase in side stickiness *κ*_*side*_ (below 75% HDI), whereas in OCD there was additionally a mild reduction in side stickiness (Kanen et al. 2019). In other words, the computational profile of GD and OCD was distinct. Perseveration is not a unitary construct (see also (Sandson and Albert 1984)): side stickiness may be representative of motor perseveration, whereas stimulus stickiness reflects stimulus perseveration. Side stickiness may therefore represent excessive motor perseveration. In contrast, the reduction in stimulus stickiness may reflect another form of behavioral inflexibility that is overly exploratory yet outcome insensitive. Low stimulus stickiness in GD detected during trial and error learning in a laboratory setting may therefore reflect a real-life increase in exploration of choices in an attempt to identify an optimal strategy, e.g., tracking new stimuli in a casino game (Clark 2010). Once the ‘optimal’ stimulus has been chosen following exploration, greater side stickiness may result in motor perseveration resulting in excessive losses. Whilst one interpretation of low stimulus stickiness in OCD is that it is a manifestation of increased checking behavior, this too can be thought of as maladaptive exploration albeit pertinent to a different real-life setting (Hauser et al. 2017; Kanen et al. 2019). Exploration (particularly of stimuli) is presumably meant to collect information; however, when such behavior becomes disconnected from outcomes it may contribute to compulsions in GD and OCD. It remains to be determined how the neural mechanisms supporting low stimulus stickiness in GD and OCD differ or overlap. Overall, value-free contributors to choice behavior have allowed for novel dissociations of GD, OCD, and SUD, and point to possible computational fingerprinting, which could eventually be useful for informing psychiatric classification.

At the neural level, group differences were also observed during ongoing RL processes. Differences in brain activity when tracking reward and punishment EVs were seen in participants with GD. In these individuals, there was greater activity in response to reward EVs in the amygdala, hippocampus and cingulate gyrus compared to HCs. When tracking punishment EV, on the other hand, there was lower activity in the postcentral gyrus, superior parietal lobule and occipital areas, suggesting that individuals with GD differentially track EVs of stimuli in their surroundings in favor of reward-related expectancies. In the CUD group, there was also an altered balance in RL, instead with lower responses to positive PPEs and greater responses to negative PPEs in the SFG and neighboring regions compared to control participants, which suggests preferential processing of punishment. This aligns with our recent finding that individuals with SUD show increased punishment learning rates (Kanen et al. 2019). In summary, there appear to be uniquely aberrant neural signals in each patient group when tracking value-related information important for RL processes.

By linking the computational modeling parameters to the fMRI data, we also identified regions involved in the modulation of RL measures, which has not been investigated in previous human studies. We found that the learning rate parameter for reward (*α*_*rew*_) was correlated with areas that responded to RPEs and PPEs, including the SFG, MFG, cingulate and paracingulate gyri. Therefore, these regions appear to be of key importance for RL and are likely to be involved in the modulation of the reward learning rate (*α*_*rew*_). The SFG and ACC are key areas underlying error and action monitoring, providing support for their involvement in reward learning (Carter et al. 1998; Botvinick et al. 1999). Moreover, a meta-analysis including 35 studies reported that these areas are consistently activated when there is a prediction error (Garrison et al. 2013).

At least two previous studies have reported reduced learning rates, reinforcement sensitivity and increased stimulus stickiness in individuals with SUD compared with HCs (Kanen et al. 2019; Lim et al. 2021). In the present study, meanwhile, we observed diminished reward learning rates and increased stimulus stickiness in CUD only when contrasted with GD. Duration of substance abuse may be a key factor underlying the less pronounced RL results in CUD when compared to these two previous studies. Whereas the CUD sample in the present study had an average duration of substance use of 3.7 years (Verdejo-Garcia et al. 2015), the participants with SUD in previous studies reporting more pronounced RL deficits had been using for an average of 11.7 (Ersche et al. 2011) and 13.7 years (Lim et al. 2021). Additionally, a criterion in our study was abstinence, which was not the case in the other two investigations. These differences in sample suggest longer exposure to substances may have more pronounced effects on RL processes, possibly due to neurotoxicity, and may therefore help reconcile the RL findings between these studies. As GD itself does not involve substance use, we would not expect the same magnitude or mechanism of change in RL effects related to disease duration. At the same time, such contrasts between GD and SUD may inform which aspects of RL in SUD are more or less likely to be tied to neurotoxic effects.

Another consideration when reconciling this series of studies is differences in study design. Lim and colleagues, for example, used a probabilistic task which had separate conditions for reward and punishment, tested in individuals with CUD (Lim et al. 2021), while Kanen et al. included individuals with any type of SUD (Kanen et al. 2019) – this may also explain the decreased punishment learning rate observed in the former compared to an increased punishment rate in the latter. It has been shown that individuals with Cocaine and Amphetamine Use Disorders perform differently during reversal learning, with increased perseveration seen preferentially in those with CUD (Ersche et al. 2008). This difference may relate to elevated stimulus stickiness in CUD observed here only when contrasted with GD.

Based on the neural results presented here, individuals with GD appear to be less sensitive to punishment EV but more sensitive to reward EV than controls. A study of performance on a two-choice lottery task found that choice behavior in GD patients was less sensitive to EVs for both reward and punishment, with this group using information about magnitude and probability information less than HCs (Limbrick-Oldfield et al. 2021). Thus, attenuated responses to punishment appear to be common across tasks in GD. Although sensitivity to reward was increased in our study and decreased in Limbrick-Oldfield et al. (2021), this may have been because different behavioral paradigms were used. Consistent with our findings, a previous study employing a card-guessing task, participants with GD had increased neural responses in the VS and OFC when tracking reward EV (Van Holst et al. 2012). Overall, these studies suggest that GD patients show altered responses to reinforcement tracking and are less sensitive to punishment.

In individuals with SUD, reduced responses to PEs in the VS and mOFC on the IGT have been reported previously (Tanabe et al. 2013). In a separate study using electroencephalography, impaired RPE signaling in CUD was also found (Parvaz et al. 2015). In contrast, we found increases in responding to PPEs, rather than reduction in RPEs. Following cocaine abstinence in individuals with CUD, enhanced signals to positive PEs, regardless of whether reward or punishment was predicted, have been observed (Wang et al. 2019). Although we report reduced activity following positive PPEs, this may be because we separated reward and punishment PEs and suggests that the two PEs are differentially altered in CUD. Altered responses to PE related to both reward and punishment could be a contributor to compulsive drug use, as it persists despite negative outcomes. In patients with OCD, RPE responses were altered in the nucleus accumbens and anterior cingulate cortex, further highlighting that RL can be used to distinguish disorders of compulsivity, both through behavior and its associated neural substrates (Murray et al. 2019).

We report that stimulus stickiness (^κ^*stim*) was positively correlated with activity in the dorsolateral PFC (dlPFC) and ventrolateral PFC (vlPFC), areas important for cognitive control, including conflict monitoring and motor inhibition, respectively (Badre and Wagner 2004; Levy and Wagner 2011). In the results presented here, patients with GD and CUD showed a stronger positive correlation with stimulus stickiness (^κ^*stim*) in these regions. This result was contrary to our expectations and previous studies, as it was predicted that stickiness would be related to *reduced* activity in these regions. A possible interpretation of this finding is that stimulus stickiness reflects bias towards one of the presented stimuli, ideally the majority reinforced one, and that the MFG and IFG are active in order to overcome this response following a reversal. However, this hypothesis would need to be explored further in future studies.

It has been demonstrated previously that both the dlPFC and vlPFC are affected in GD and CUD (Goldstein and Volkow 2011; Raimo et al. 2021); here we provide a novel computational mechanism pertinent to compulsions that is linked to these regions in GD and CUD. Previous studies have demonstrated that response shifting on the PRL task is associated with vlPFC activation in control participants (Cools et al. 2002). Consistent with the present results, a prior analysis of this dataset showed the vlPFC was engaged during response shifting, yet both clinical groups showed lower vlPFC activity than HCs (Verdejo-Garcia et al. 2015). Reduced vlPFC activity during shifting has been also reported in OCD patients (Remijnse et al. 2006). These findings from previous studies, however, focus on response shifting on certain trials, whereas our analysis investigated stickiness across all trials, reflecting an overall tendency. Additionally, stickiness represents repeated responses, rather than response shifts. In rats, it has been shown that side stickiness (stimulus stickiness was not studied) is correlated with activity in medial PFC and dorsal striatal regions (Zühlsdorff et al. 2023). It is therefore possible that side and stimulus stickiness recruit different neural circuits, but this requires further analysis in the same species.

In summary, we provide novel behavioral and neural insights into GD through computational modeling of RL processes. Critically, we demonstrate that individuals with GD and CUD display perseverative behavior during PRL that differs both qualitatively and quantitatively, advancing the notion that compulsivity is not a unitary construct. We also provide evidence that individuals with GD and CUD display aberrant and opposing neural responses to rewards and punishments, in relation to expected value and PPEs. Furthermore, we link RL parameters to regions that may be involved in their modulation, which has not previously been investigated in the human literature, such as the finding that stimulus stickiness is positively correlated with activity in the dlPFC and vlPFC, areas involved in modulating the balance between goal-directed and habitual behaviors. We demonstrate that RL modeling combined with fMRI may provide new insights into the mechanisms underlying compulsive disorders and therefore refine our understanding of compulsivity transdiagnostically.

## Funding

The collection of data for this study was originally funded by the Spanish Ministry of Health / Plan Nacional Sobre Drogas and the Centre for Gambling Research at the University of British Columbia (UBC). R.N.C.’s research is supported by the UK Medical Research Council (MRC) (MR/W014386/1). K.Z. was supported by the Institute for Neuroscience at the University of Cambridge, the Alan Turing Institute, London and the Angharad Dodds John Bursary in Mental Health and Neuropsychiatry, Downing College, Cambridge. J.W.D. has received funding from GlaxoSmithKline and Boehringer Ingelheim Pharma GmbH and is a co-investigator on an MRC program grant (MR/N02530X/1). T.W.R. is also a co-investigator of the latter grant. J.W.K. was supported by an Angharad Dodds John Bursary in Mental Health and Neuropsychiatry. A.V.G is supported by a Leadership Investigator Grant from the Australian National Health and Medical Research Council (GNT2009464).

## Declaration of Interests

The Centre for Gambling Research at UBC (L.C.) is funded by support from the Province of British Columbia and the British Columbia Lottery Corporation. J.W.D. has received research grants from Boehringer Ingelheim Pharma GmbH and GlaxoSmithKline and receives royalties from Springer Verlag. T.W.R. discloses consultancy with Cambridge Cognition; he receives editorial honoraria from Springer-Nature and Elsevier and a research grant from Shionogi. R.N.C. consults for Campden Instruments and receives royalties from Cambridge Enterprise, Routledge, and Cambridge University Press.

