## Supplementary material for "Computational modeling of reinforcement learning and functional neuroimaging of probabilistic reversal dissociates compulsive behaviors in Gambling and Cocaine Use Disorders": Supplementary figure legends.docx

**SF 1.** Feedback/cue presentation – differences between healthy controls and participants with

GD (MNI coordinates: X=31, Y=-68, Z=27). Activity was higher in the GD group in the

indicated areas. Colour bar on the right-hand side represents t-statistic.

**SF 2.** Feedback/cue presentation – differences between healthy controls and participants with

GD (MNI coordinates: X=39, Y=34, Z=17). Activity was higher in the CUD group in the

indicated areas. Colour bar on the right-hand side represents t-statistic.

**SF 3.** Areas that have a stronger negative correlation with α_rew_ in the GD group than in

healthy controls during reward EV tracking (MNI coordinates: X=-32, Y=12, Z=52). Colour

bar on the right-hand side represents t-statistic.

**SF 4.** Areas that have a stronger negative correlation with α_rew_ in the CUD group than in

healthy controls when responding to positive PPE (MNI coordinates: X=-5, Y=37, Z=22).

Colour bar on the right-hand side represents t-statistic.
