## Supplementary figures and images for "Computational modeling of reinforcement learning and functional neuroimaging of probabilistic reversal dissociates compulsive behaviors in Gambling and Cocaine Use Disorders"

### SF.1.png

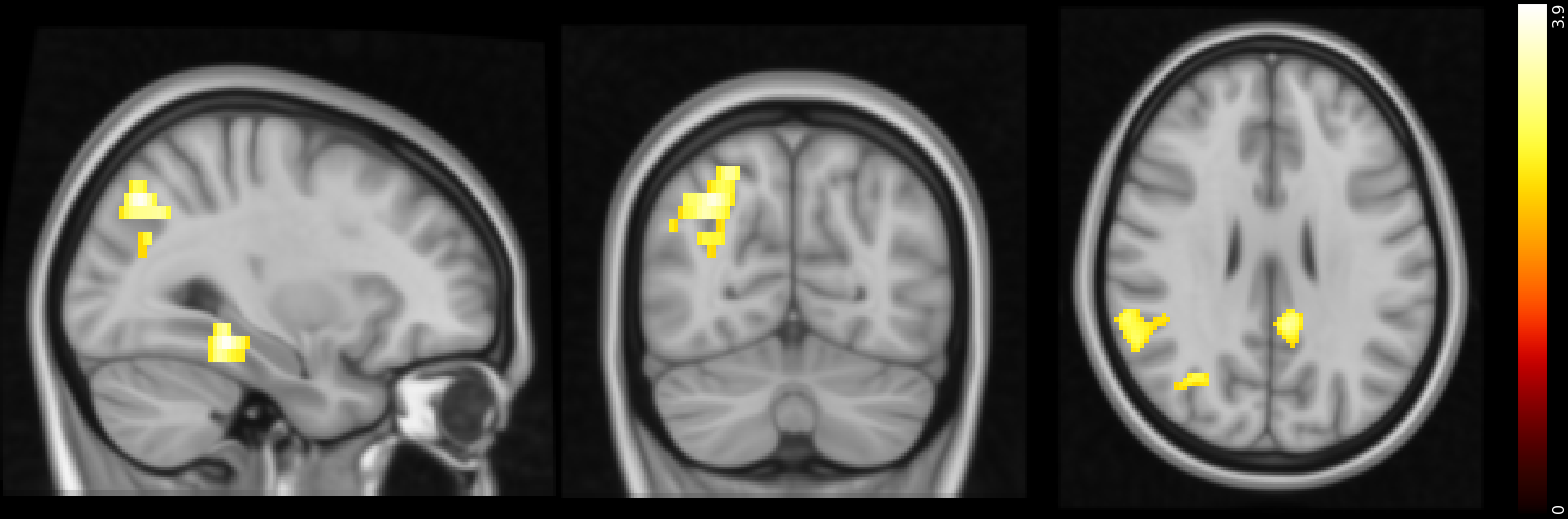

### SF.2.png

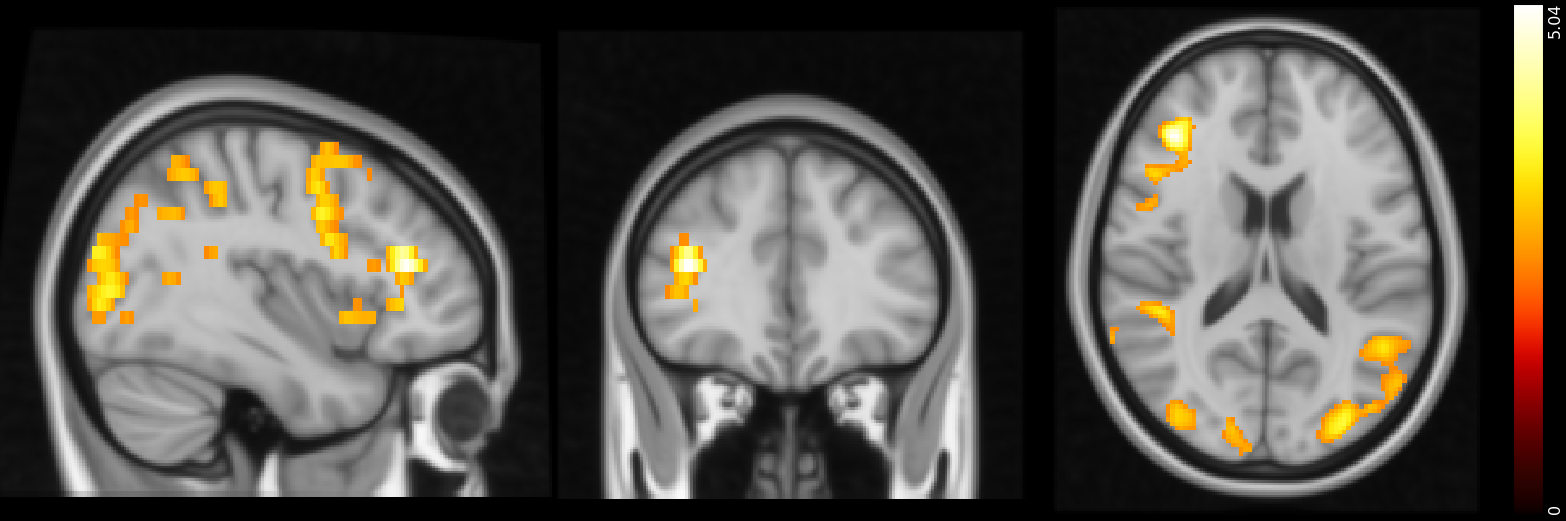

### SF.3.png

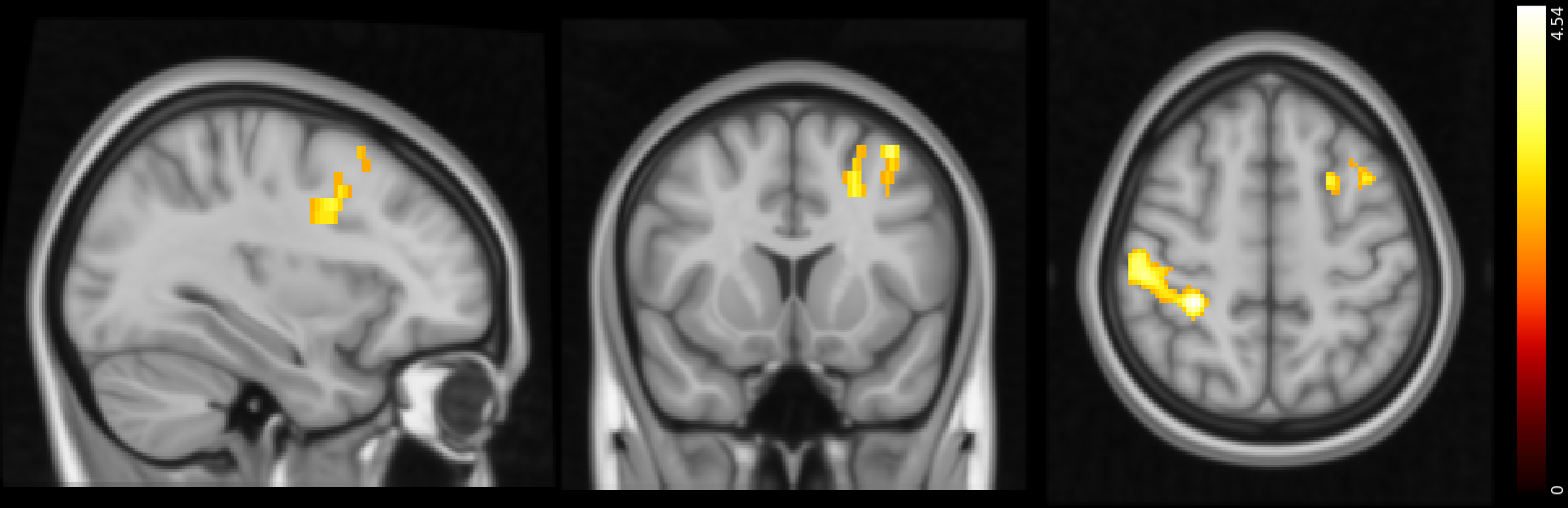

### SF.4.png

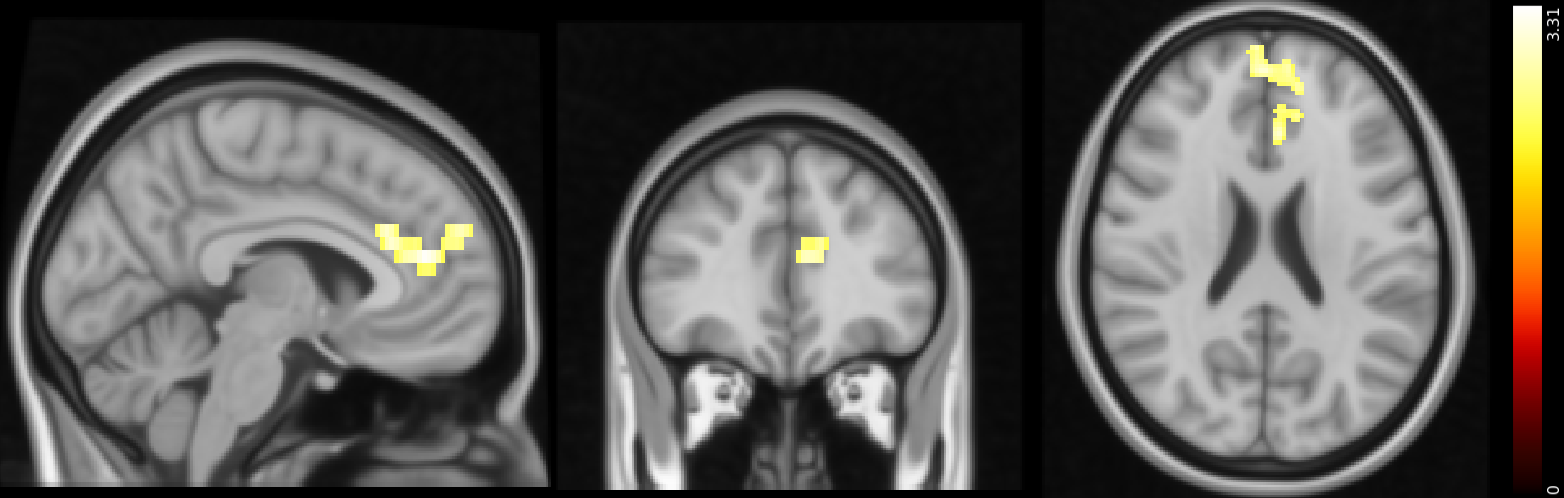
